# Integrin-dependent YAP signaling requires LAMTOR1 mediated delivery of Src to the plasma membrane

**DOI:** 10.1101/585349

**Authors:** Marc R. Block, Molly Brunner, Théo Ziegelmeyer, Dominique Lallemand, Mylène Pezet, Genevieve Chevalier, Philippe Rondé, Bernhard Wehrle-Haller, Daniel Bouvard

## Abstract

YAP signaling has emerged as an important signaling pathway involved in several normal and pathological processes. While main upstream effectors regulating its activity have been extensively studied, the interplay with other cellular processes has been far less analyzed. Here, we identified the LAMTOR complex as a new important regulator of YAP signaling. We uncovered that p18/LAMTOR1 is required for the recycling of Src on late endosomes to the cell periphery, and consequently to activate a signaling cascade that eventually controls YAP nuclear shuttling. Moreover, p18/LAMTOR1 positives late endosomes distribution is controlled by β1 integrins, extracellular matrix stiffness and cell contractility. This likely relies on the targeting of microtubules to β1 positive focal adhesion via ILK. Altogether our findings identify the late endosomal recycling pathway as a major regulator of YAP.

## Introduction

Cellular and developmental processes are tightly regulated by external inputs. Amongst these stimuli, extracellular factors such as transforming growth factor-β (TGF-β), bone morphogenetic proteins (BMPs), or serum components activate their respective receptors at the cell surface to initiate intracellular signaling. However, these signaling systems require an integrin-dependent cell-matrix adhesion to be effective [1,2]. Indeed, over the last decade, the physical and chemical properties of the extracellular matrix have emerged as major players that tune those factor-dependent signaling pathways [3,4]. This regulation by the extracellular environment acts at various levels such as the adhesion-regulated formation of signaling platforms at the cell surface [5], the stimulation of endocytosis and vesicular trafficking [6–8], or the control of nuclear shuttling of specific transcription factors and chromatin remodelers to modulate gene expression [3,9,10].

Among the mechano-sensitive pathway, the two related transcriptional cofactors YAP (yes associated protein) and TAZ (transcriptional co-activator with PDZ-binding motif) emerged as crucial players in this response. Indeed, the nuclear shuttling of YAP and TAZ is under the direct control of the compliance and composition of the extracellular matrix, as well as the cell shape and cell confluence [11]. In the nucleus YAP and TAZ, interact with members of the TEAD family to form functional transcription factors and drive or modulate gene expression [9]. In turn, these expression profiles will affect cell behavior such as proliferation/differentiation or migration, and thereby enable the integration of external factors and extracellular matrix-dependent cues for cells to adapt to their extracellular environment [12]. Over the last decade, the core Hippo signaling pathway leading to YAP and TAZ nuclear translocation has been extensively studied giving important insights in cell physiology and organ growth. However, how the mechanical cues embedded within the extracellular matrix are translated into a functional regulation of YAP and TAZ nuclear shuttling remains elusive. We and others have identified that integrin-dependent cell adhesion and in particular β1 integrins are of critical importance for YAP/TAZ nuclear shuttling [13,14]. Indeed, β1 integrin dependent cell adhesion-signaling, likely through the Src family kinases, leads to the recruitment and activation of Rac-1 at protrusive cell borders, and stimulates a PAK1 dependent signaling cascade resulting in merlin phosphorylation. Phosphorylated merlin releases YAP from a merlin/LATS/YAP complex and thereby permits its nuclear translocation [14]. While, the role of Rac-1 and Src in YAP nuclear translocation have been reported by several groups, the mechanistic involvement of integrins in this process remained unexplored [15–17]. Here in, we discovered a functional link between β1-integrin-dependent recycling and the involvement of vesicular trafficking in the regulation of YAP nuclear translocation. Based on a genetic screen, we identified the long recycling loop as an important process in regulating YAP nuclear shuttling. Amongst the vesicles that recycle back to the plasma membrane, the late endosomal compartment (containing the so-called “detergent resistant membrane” DRM) appeared of paramount importance. The loss of LAMTOR1, a protein essential for docking the LAMTOR complex to late endosomal DRMs, severely affected YAP nuclear shuttling, as well as late endosomes distribution. As observed for YAP activation and LAMTOR localization, the late endosome subcellular distribution was found to be cell adhesion dependent and mechanosensitive. This latter distribution of the endosomal compartment relies on a functional microtubule network organized by β1 integrins through ILK. Mechanistically, these data highlight a new role for the LAMTOR complex in Src recycling back to the plasma membrane to control YAP nuclear shuttling.

## Results

### Vesicular recycling to the plasma membrane regulates YAP nuclear translocation

While the core signaling pathway leading to YAP nuclear translocation has been extensively studied over the last years [18,19], the role of vesicular trafficking has not been analyzed. To address the role of vesicular trafficking in YAP localization, we used a genetic screen aimed at blocking vesicular trafficking at different steps. Expression of Rab dominant negative mutants is a common way to investigate the involvement of cellular trafficking in a variety of cellular processes [20]. Dominant negative constructs of Rab-4, −5, −7 and −11, were selected to target the short recycling loop, early endosomes, the late endosomal/endo-lysosomal compartment, or the long recycling loop, respectively. Dominant negative, as well as their wild-type forms (controls), were transiently expressed in mesenchymal osteoblastic cells and YAP subcellular localization was assayed by immunostaining. Under each experimental condition, the relative nuclear to cytoplasmic ratio of YAP was quantified using confocal imaging. Neither blocking the maturation of late endosomes into endo-lysosomes with Rab-7^T22N^, nor inhibiting the short recycling loop with Rab-4^S22N^ significantly affected YAP nuclear localization (Fig. 1A and EV1). In contrast, the expression of Rab-5^S34N^ or Rab-11^S24N^ induced a significant negative effect on YAP nuclear localization (Fig. 1A and Fig. EV1). These data suggested that endocytosis and the long recycling loop might be involved in YAP nuclear translocation.

**Figure 1.**
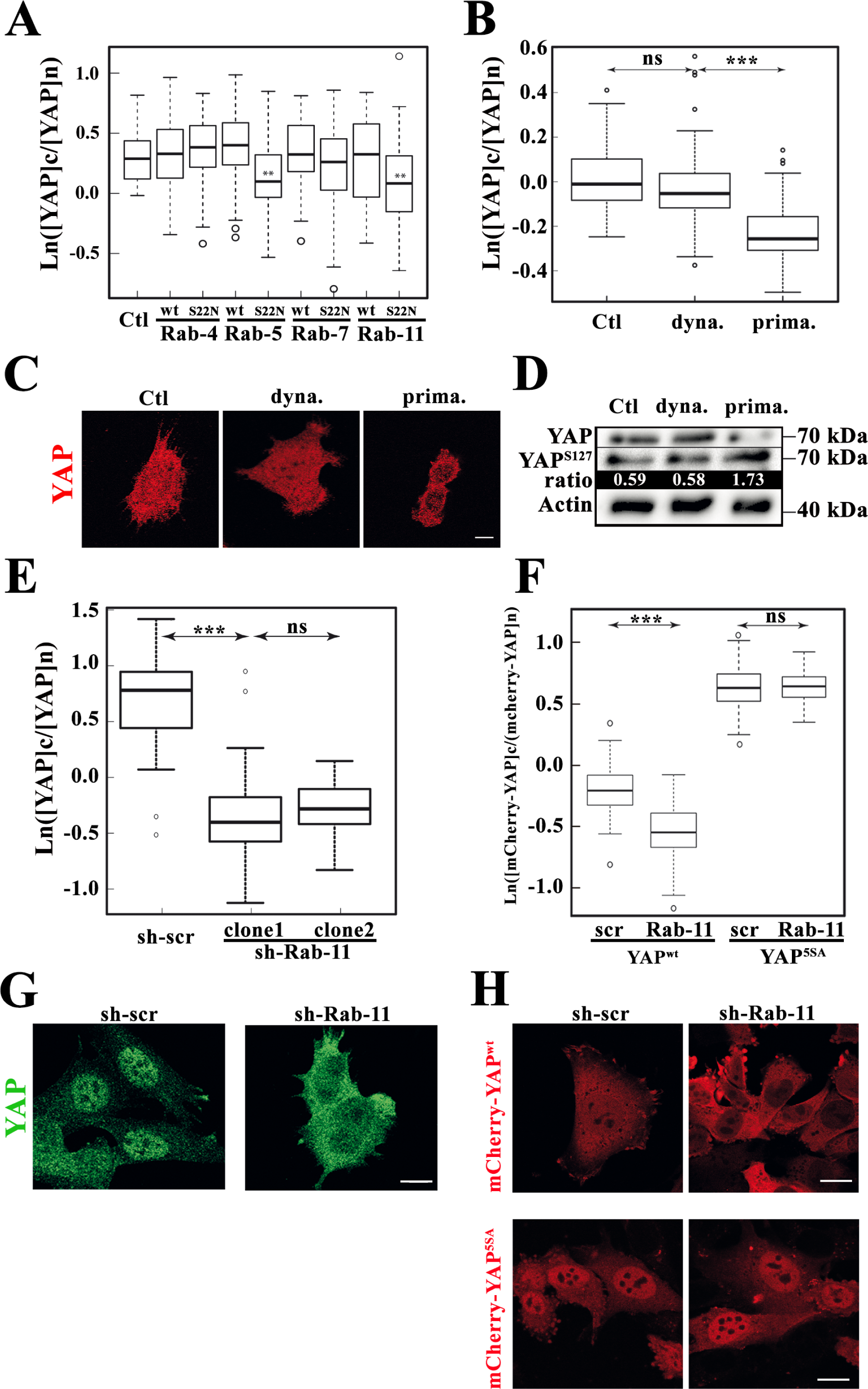
Rab-11-dependent vesicular recycling control YAP nuclear shuttling. **A.** Statistical analysis of YAP cytoplasmic to nuclear ratio represented in a logarithmic scale. Mesenchymal osteoblast cells (β1^f/f^) were transiently transfected with either wild-type or dominant negative form of GFP-Rab fusion proteins. 24h post transfection cells were seeded on glass coverslips and YAP subcellular localization analyzed by indirect immunofluorescence. Intensity values were measured from confocal images using Fiji software. p-values were assessed by a two-tailed unpaired Student’s t-test, the box plot is representative of 2 independent experiments with n>50 cells analyzed. **B.** Statistical analysis of YAP cytoplasmic to nuclear ratio represented in a logarithmic scale. Osteoblast cells (β1^f/f^) were seeded overnight on glass coverslips and either dynasore, 10µg/ml, or primaquine, 50µg/ml, were added to the culture medium for 1h. YAP subcellular localization was then analyzed by indirect immunofluorescence. Intensity values were measured from confocal images using Fiji software. p-values were assessed by a two-tailed unpaired Student’s t-test, the box plot is representative of 3 independent experiments with n>30 cells analyzed. **C.** Immunostaining of YAP (red) in osteoblast cells seeded overnight on glass coverslips and treated with or without dynasore, 10µg/ml or primaquine, 50µg/ml, respectively. Scale bar represents 10µm. **D.** Western blot analysis of YAP, and YAP^pS127^, in osteoblast cells treated with or without dynasore, 10µg/ml or primaquine, 50µg/ml as described above. Labeling was estimated with Chemidoc CCD camera (Biorad) and ratios were estimated using Image Lab software (Biorad). Actin was used as internal loading control. Representative of 3 independent experiments. **E.** Statistical analysis of YAP cytoplasmic to nuclear ratio represented in a logarithmic scale. Osteoblast cells (β1^f/f^) expressing sh-scramble (sh-scr) or sh-Rab-11a were seeded overnight on glass coverslips. YAP subcellular localization was then analyzed by indirect immunofluorescence. Intensity values were measured from confocal images using Fiji software. p-value was assessed by a two-tailed unpaired Student’s t-test, the box plot is representative of 2 independent experiments with n>30 cells analyzed. **F.** Statistical analysis of YAP cytoplasmic to nuclear ratio represented in a logarithmic scale. Osteoblast cells (β1^f/f^) bearing stable sh-scramble (scr) or sh-Rab-11a (Rab-11) together with mCherry-YAP^wt^ or mCherry-YAP^5SA^ were seeded overnight on glass coverslips. YAP subcellular localization was then analyzed by imaging mCherry fluorescence with a confocal microscope. Intensity value was measured using Fiji software. p-value was assessed by a two-tailed unpaired Student’s t-test, the box plot is representative of 2 independent experiments with n>30 cells analyzed. **G.** Immunostaining of YAP (green) in osteoblast cells stably expressing sh-scramble (scr) or sh-Rab-11a (Rab-11) and seeded overnight on glass coverslips. Scale bar represents 10µm. **H.** Subcellular localization of mCherry-YAP^wt^ or mCherry-YAP^5SA^ (red) in osteoblast cells stably expressing sh-scramble (scr) or sh-Rab-11a (Rab-11). Scale bar represents 10µm.

Next, genetic data were complemented with pharmacological inhibitions using well-established vesicular trafficking inhibitors such as dynasore (that inhibits dynamin dependent endocytosis), and primaquine (PQ, that inhibits both the long and short recycling loop) [21,22]. During the time course of the experiment in fully spread cells, dynasore treatment failed to significantly perturb YAP nuclear localization when compared to control cells (Fig. 1B and 1C). These results apparently did not fit with the above results obtained with Rab-5^S34N^ and shed some doubts on the role of endocytosis in controlling YAP nuclear translocation. Conversely, the addition of primaquine was consistent with the data obtained with dominant negative data, resulting in an important reduction of YAP nuclear localization when compare to control cells (Fig. 1B and 1C). In addition, primaquine treatment increased the fraction of phosphorylated YAP versus total YAP when compared to control cells, while dynasore-treated cells did not show significant effects on YAP-phosphorylation (Fig. 1D). Altogether these data suggested that the long recycling loop plays a role in the dephosphorylation of YAP and the regulation of YAP nuclear translocation, while the role of endocytosis could not be clearly established.

In the light of these data, we further focused our investigations on the long recycling loop. It is well-known that the over-expression of small GTPases might have unwanted side effects by titrating important effectors shared by other signaling pathways. To circumvent this issue, we generated stable cell lines in which Rab-11a expression was reduced by sh-RNA mediated silencing. YAP subcellular localization was then analyzed and quantified by immunofluorescence. Silencing of Rab-11a reduced both the nuclear localization of endogenous YAP, and that of stably expressed mCherry-YAP fusion protein (Fig. 1E-1G). In contrast, the expression of the non-phosphorylatable form of YAP (mCherry-YAP^5SA^) was insensitive to Rab-11 silencing (Fig 1H), suggesting that a Rab-11 stimulated process mediated YAP-dephosphorylation and nuclear translocation.

### p18/LAMTOR1 controls YAP nuclear translocation

We, and others, previously reported that cell adhesion plays an important role in regulating YAP nuclear translocation [14,15,23]. It is noteworthy that cell adhesion is also implicated in detergent resistant membrane (DRM) trafficking back to the plasma membrane [24,25]. Knowing the role of DRM in cell signaling, we wondered whether the requirement of the long recycling loop for YAP nuclear translocation could involve DRM recycling and its associated signaling. We focused our attention on LAMTOR since this complex has been shown to be recruited onto DRMs and its genetic deletion is associated with reduced proliferative capabilities [26,27].

DRMs were labelled using p18/LAMTOR1-GFP and their subcellular distribution was analyzed in either Rab-11a (sh-Rab-11) or scrambled (sh-scr) silenced cells. In control cells, p18/LAMTOR1 positive vesicles were distributed throughout the cells with a clear localization of vesicles at the cell edges. However, silencing of Rab-11a led to a significant reduction in those peripheral vesicles. Conversely, perinuclear vesicles were not affected by the loss of Rab-11a (Fig. 2A and 2C). We also noticed a reduced number of punctate staining in the whole cytoplasm with an average of 381+/-148 and 206+/-109 vesicles per cell in sh-scramble and sh-Rab-11a, respectively (Fig. 2A). These data were further supported by video-microscopy that revealed an important defect of p18/LAMTOR1 vesicle velocity compared to controls (Movies EV1 and EV2). In line with these results, blocking vesicular recycling with primaquine led to a massive redistribution of p18/LAMTOR1 toward the perinuclear region together with a depletion of the peripherally located pool of vesicles (Fig 2B and 2C).

**Figure 2.**
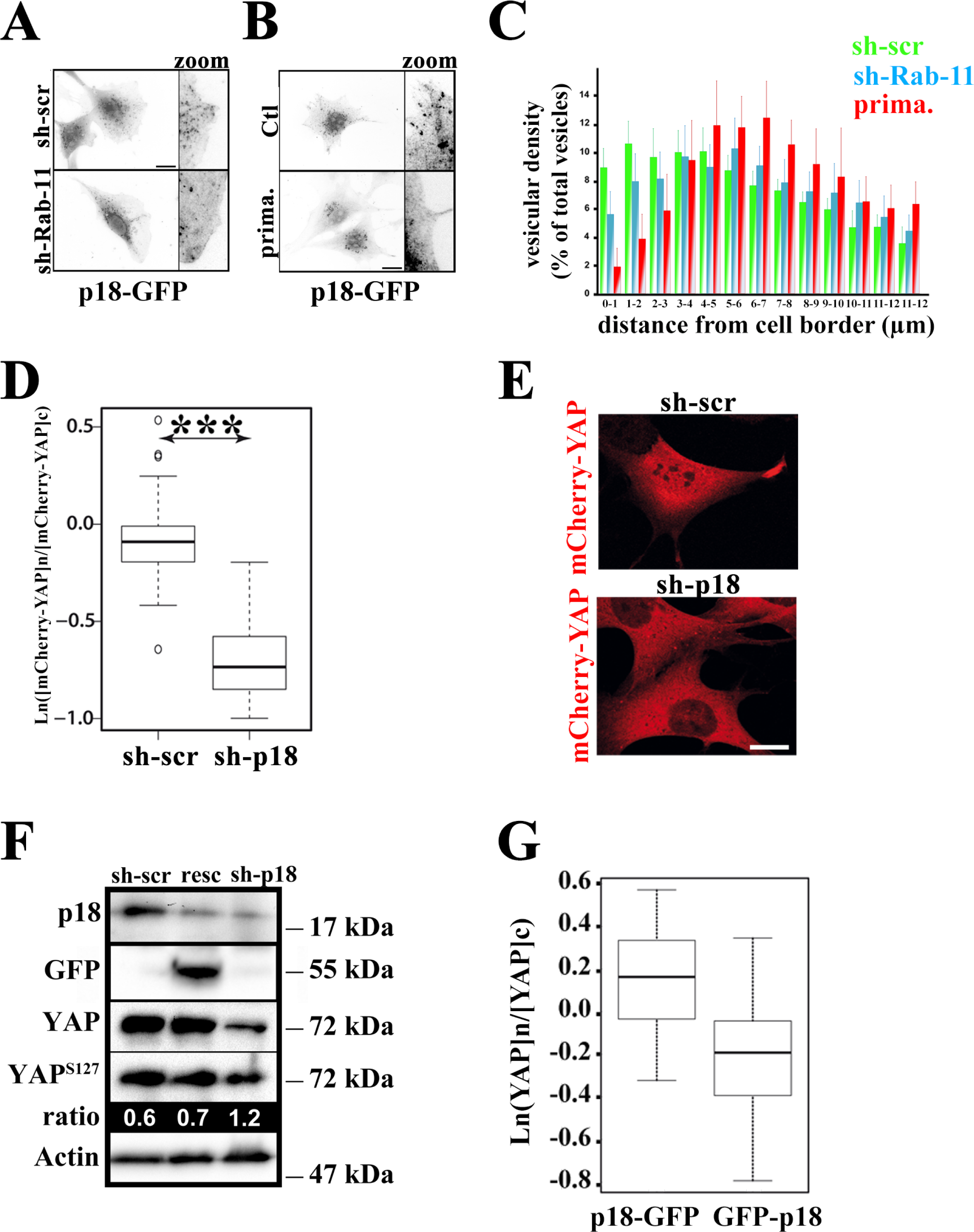
p18/LAMTOR1 distribution is Rab-11-dependent and controls YAP nuclear shuttling. **A.** p18/LAMTOR1 subcellular distribution in sh-scr or sh-Rab-11a expressing osteoblast cells. Scale bar represents 10µm. **B.** p18/LAMTOR1 subcellular distribution in osteoblast cells treated or not with primaquine (1h, 50µg/ml). Scale bar represents 10µm **C.** Statistical analysis of p18 -GFP subcellular distribution. Osteoblast cells stably expressing either sh-scr (red), stable sh-Rab-11a (blue) or treated with primaquine (green, 1h, 50 µg/ml), were imaged. Fluorescence distribution of p18 -GFP was analyzed using Icy software. Histograms represent stepwise particles localization from the cell edges to the cell nucleus displayed as percent of total vesicles. Representative of 2 independent experiments. **D.** Statistical analysis of YAP cytoplasmic to nuclear ratio represented in a logarithmic scale. Osteoblast cells stably expressing mCherry-YAP^wt^ and bearing sh-scramble (sh-scr) or sh-p18/LAMTOR1 were seeded overnight on glass coverslips. YAP subcellular localization was then analyzed by imaging mCherry fluorescence with a confocal microscope. Intensity values were obtained using Fiji software. p-value was assessed by a two-tailed unpaired Student’s t-test, the box plot is representative of 2 independent experiments with n>30 cells analyzed. **E.** Subcellular localization of mCherry-YAP^wt^ (red) in osteoblast cells expressing stable sh-scramble (scr) or sh-p18/LAMTOR1 (sh-p18). Scale bar represents 10µm. **F.** Western blot analysis of YAP, YAP^pS127^ and p18/LAMTOR1 expression in osteoblast cells. Labeling was estimated with Chemidoc CCD camera (Biorad) and ratios were estimated using Image Lab software (Biorad). Actin was used as internal loading control. Representative of 3 independent experiments. **G.** Statistical analysis of YAP cytoplasmic to nuclear ratio represented in a logarithmic scale. Osteoblast cells (β1^f/f^) expressing either p18 -GFP or the dominant negative form GFP-p18 were seeded overnight on glass coverslips. YAP subcellular localization was then analyzed by indirect immunofluorescence. Intensity values were obtained from confocal images using Fiji software. p-value was assessed by a two-tailed unpaired Student’s t-test, the box plot is representative of 2 independent experiments with n>30 cells analyzed.

To directly address the role of LAMTOR1 in YAP signaling, cells with a stable sh-RNA directed against p18/LAMTOR1 (sh-p18) were generated. In those cells, a significant reduction in YAP nuclear staining was observed when compared to sh-scramble (Fig. 2D and 2E). In good agreement with these data, YAP was hyperphosphorylated in p18 silenced cells when compared to sh-scramble or rescued cells (Fig 2F). We also observed a strong decrease in total YAP level likely reflecting its degradation via the previously described phosphodegredon [28]. The anchoring of p18/LAMTOR1 to DRM requires some lipid modifications occurring at its N-terminal part. Therefore, the generation of an N-terminally fused GFP is predicted to infer with its function and recruitment onto DRM surfaces. Indeed, in sharp contrast to p18/LAMTOR1-GFP that localizes to vesicles, the distribution of n-terminally tagged GFP-p18/LAMTOR1 was diffuse within the cells (Fig EV2). Accordingly, YAP nuclear staining was also reduced in GFP-p18/LAMTOR1 expressing cells (Fig. 2G).

### p18/LAMTOR1 subcellular distribution is β1 integrin dependent and mechanosensitive

Since our data showed that p18/LAMTOR1 is involved in the nuclear shuttling of YAP, we wondered whether exposing cells to experimental conditions known to modulate YAP nuclear localization might also affect p18/LAMTOR1 distribution. Amongst the numerous inputs that modulate YAP nuclear localization, cell adhesion, matrix stiffness and cell contractility are of importance and altogether account for the mechano-dependent cellular response of YAP [10,11,19,28]. However, whether mechanical cues also affect LAMTOR1 or, more broadly, the late endosome distribution has never been investigated to the best of our knowledge. Cells expressing either mCherry-YAP or p18/LAMTOR1-GFP were seeded onto 2, 10, or 30 kPa fibronectin coated PDMS hydrogels for 2 hours and p18/LAMTOR1-GFP or mCherry-YAP subcellular localization was analyzed. Consistent with previous reports, cells on soft (2 kPa) fibronectin-coated PDMS hydrogels showed reduced YAP nuclear staining when compared to stiffer hydrogels (10 or 30 kPa) (Fig. 3A and 3B). Seeding cells expressing GFP tagged p18/LAMTOR1 under similar experimental conditions showed that extracellular stiffness also affected its distribution. Indeed, while cells on medium compliance or stiff fibronectin coated PDMS hydrogels (10 and 30 kPa, respectively) displayed peri-nuclear and peripheral p18/LAMTOR1 positive vesicles. Culturing cells on low stiffness (2 kPa) substrate led to a significant loss of peripherally located p18/LAMTOR1 positive vesicles (Fig. 3C and 3D). Similarly, lowering cell contractility by the addition of blebbistatin or the removal of the main mechano-transducers, i.e. β1 integrins, reduced p18/LAMTOR1 vesicular density at the cell periphery (Fig 3E and 3F). These data revealed that LAMTOR1 subcellular distribution, and that of recycling vesicles are controlled by mechanical cues and cell contractility. Altogether, these data supported the idea that at least some of DRM recycling to the plasma membrane via the LAMTOR complex is controlled by cell adhesion, contractility, and mechanical cues.

**Figure 3.**
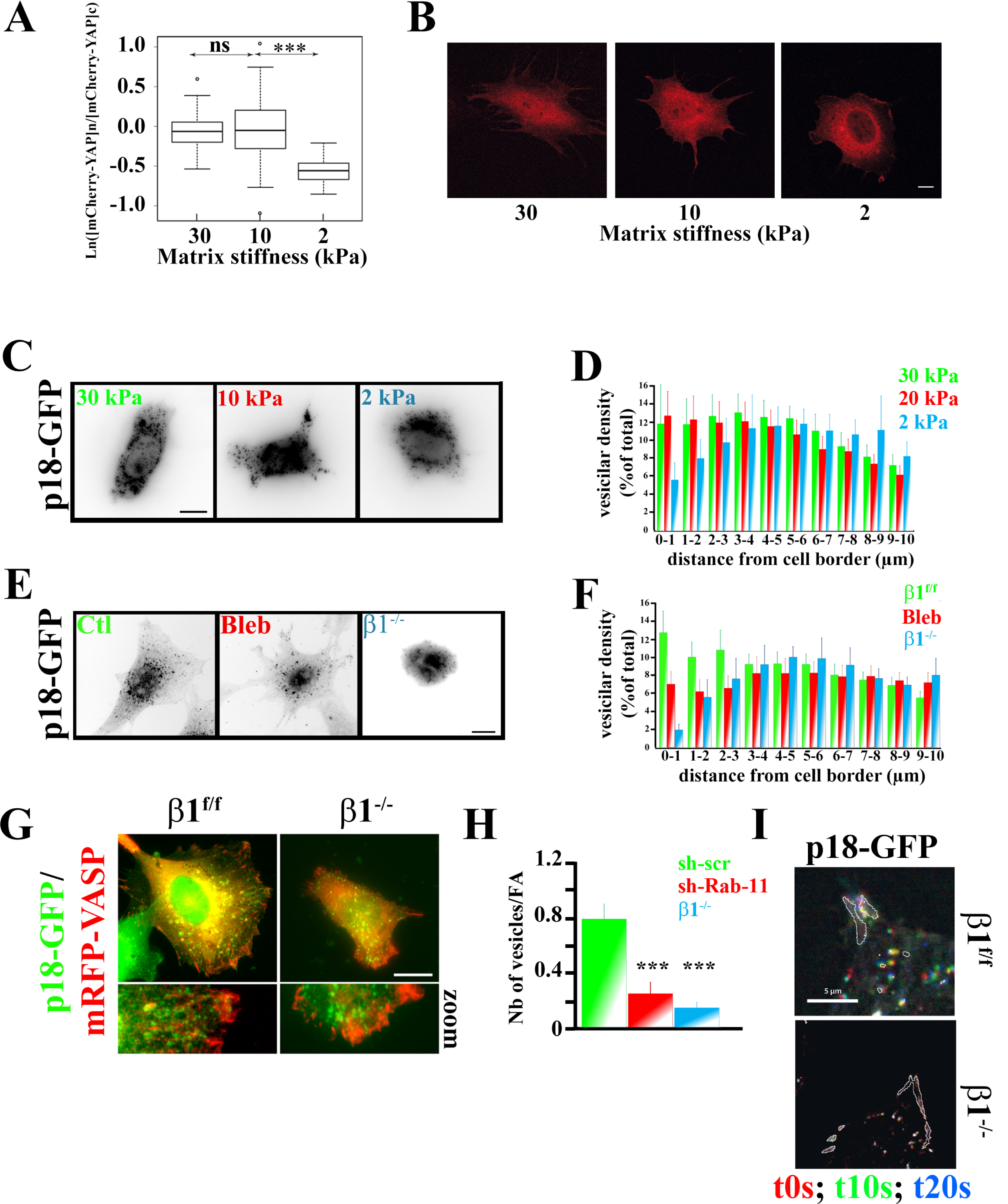
Cell adhesion, matrix stiffness and cell contractility controls p18/LAMTOR1 subcellular distribution and dynamics. **A.** Statistical analysis of YAP cytoplasmic to nuclear ratio represented in a logarithmic scale. Osteoblast cells stably expressing mCherry-YAP^wt^ were seeded for 2h on fibronectin-coated PDMS hydrogel at different stiffness (30 kPa, 10 kPa, 2kPa). YAP subcellular localization was then analyzed by imaging mCherry fluorescence with a confocal microscope. Intensity values were obtained using Fiji software. p-value was assessed by a two-tailed unpaired Student’s t-test, the box plot is representative of 2 independent experiments with n>30 cells analyzed. **B.** Subcellular localization of mCherry-YAP^wt^ (red) in osteoblast cells spread for 2h on fibronectin-coated PDMS hydrogels of different stiffness (Young’s moduli are 30 kPa, 10 kPa, 2kPa). Scale bar represents 10µm. **C.** Subcellular localization of p18/LAMTOR1 in osteoblast cells spread for 2h on fibronectin-coated PDMS hydrogel of different stiffness (Young moduli are 30 kPa, 10 kPa, 2kPa). Scale bar represents 10µm. **D.** Statistical analysis of p18/LAMTOR1-GFP subcellular distribution in osteoblast cells spread for 2h on fibronectin-coated PDMS hydrogel of different stiffness (30 kPa, 10 kPa, 2 kPa). GFP fluorescence were imaged and p18/LAMTOR1 distribution was analyzed using Icy software. Histogram represents stepwise particles localization from the cell edges to the nucleus displayed as percent of total vesicles. Representative of 2 independent experiments. p-value was assessed by a two-tailed unpaired Student’s t-test. It was non-significant except between 20 kPa and 2 kPa condition for 0-1 (***) and 1-2 (**) µm. **E.** Subcellular localization of p18/LAMTOR1. Control osteoblasts expressing p18-GFP were treated or not with blebbistatin for 1h at 10µg/ml (Bleb). β1 integrin deficient osteoblasts were seeded overnight on glass coverslips. Scale bar represents 10µm. **F.** Statistical analysis of p18/LAMTOR1-GFP subcellular distribution in untreated osteoblast (green), treated with blebbistatin (1h, 10µg/ml) or in β1 deficient cells. GFP fluorescence were imaged and p18/LAMTOR1 distribution was analyzed using Icy software. Histogram represents stepwise particles localization from the cell edges to the cell nucleus displayed as percent of total vesicles. Representative of two independent experiments. p-value was assessed by a two-tailed unpaired Student’s t-test. It was non-significant except between sh-scr and bleb for 0 to 2 (***); 2-3 (**) and 2-3 (***) µm and between sh-scr and β1^−/−^ between 0-1 (***) and 1-2 (**) µm. **G.** Control (β1^f/f^) or β1 integrin deficient osteoblasts stably expressing p18/LAMTOR1-GFP (green) and mRFP-VASP (red) were seeded overnight on glass coverslips and their relative distribution visualized. Scale bar represents 10µm. **H.** Stastistical analysis of p18/LAMTOR1 targeting to focal adhesions in control (sh-scr, green) sh-Rab-11 expressing cells (red) and β1 deficient osteoblasts (blue) (red). p-value was assessed by a two-tailed unpaired Student’s t-test, with n>20 cells analyzed. **I.** Digital analysis of p18/LAMTOR1-GFP dynamics in control cells (β1^f/f^) and β1 integrin deficient cells. Images at different time frames at 0s (red), 10s (green) and 20s (blue) were merged to visualize dynamics. Focal adhesions were visualized using mRFP-VASP. Scale bar represents 5µm.

It has been reported that a subset of p14/LAMTOR2 and Rab-7 positive vesicles are targeted to focal adhesion sites [29]; however, whether specific integrins are required for this targeting has not been addressed. Therefore, p18/LAMTOR1-GFP and mRFP-VASP were stably introduced into control and β1 integrin deficient cells that still adhered via αvβ3-dependent focal adhesions. In sharp contrast to parental cells that displayed an evident targeting of p18/LAMTOR1 to focal adhesions as previously reported, the loss of β1 integrin resulted in a significant reduction of this targeting to VASP-recruiting focal adhesions (Fig. 3G and 3H). Moreover, as evidenced by TIRF based video-microscopy, dynamics of p18/LAMTOR1 positive vesicles were severely affected by the loss of β1 integrins, as evidenced by video-microscopy, and consequently their targeting to focal adhesion sites (Movie EV3 and EV4, Fig. 3I). Similar p18/LAMTOR1 targeting defects were also observed in Rab-11 silenced cells supporting the involvement of Rab-11 dependent recycling loop in this process (Fig. 3H, Fig. EV3). Collectively, these data showed that β1 integrin dependent cell adhesion controls vesicular trafficking and focal adhesion targeting of LAMTOR positive vesicles.

### LAMTOR dynamics and localization are controlled by microtubules anchoring to Focal Adhesion and impact YAP nuclear translocation

Having shown that cell adhesion and mechanical cues are important in controlling p18/LAMTOR1 distribution, next we investigated the molecular mechanism underlying this regulation. Interestingly, cell adhesion was proposed to regulate microtubules anchoring to cell edges [25]. Taking in consideration the well-known role of microtubules in controlling vesicular dynamics we asked whether the β1 dependent distribution of p18/LAMTOR1 was involving the microtubular system. First, we analyzed the organization of the microtubule network during cell spreading on fibronectin, a well-known β1 integrin extracellular ligand. The microtubule network appeared more organized over time with an obvious increase in the number of the microtubules reaching cell borders (Fig. 4A). It is noteworthy that p18/LAMTOR1 also displayed a dynamic behavior with vesicles mainly located close to the nucleus at early spreading time that clearly dispersed during cell spreading (Fig. 4B). Therefore, the distribution of p18/LAMTOR1 positive vesicles appeared to be correlated with microtubule reorganization that normally occurs during cell spreading. In line with an essential role of microtubules, we observed that p18/LAMTOR1 positive vesicles were often associated with them (Fig. 4C, left panel), and the addition of nocodazole (to induce microtubules depolymerization) strongly redistributed p18/LAMPTOR1 at the peri-nuclear region (Fig. 4C, right panel) as previously described [29].

**Figure 4.**
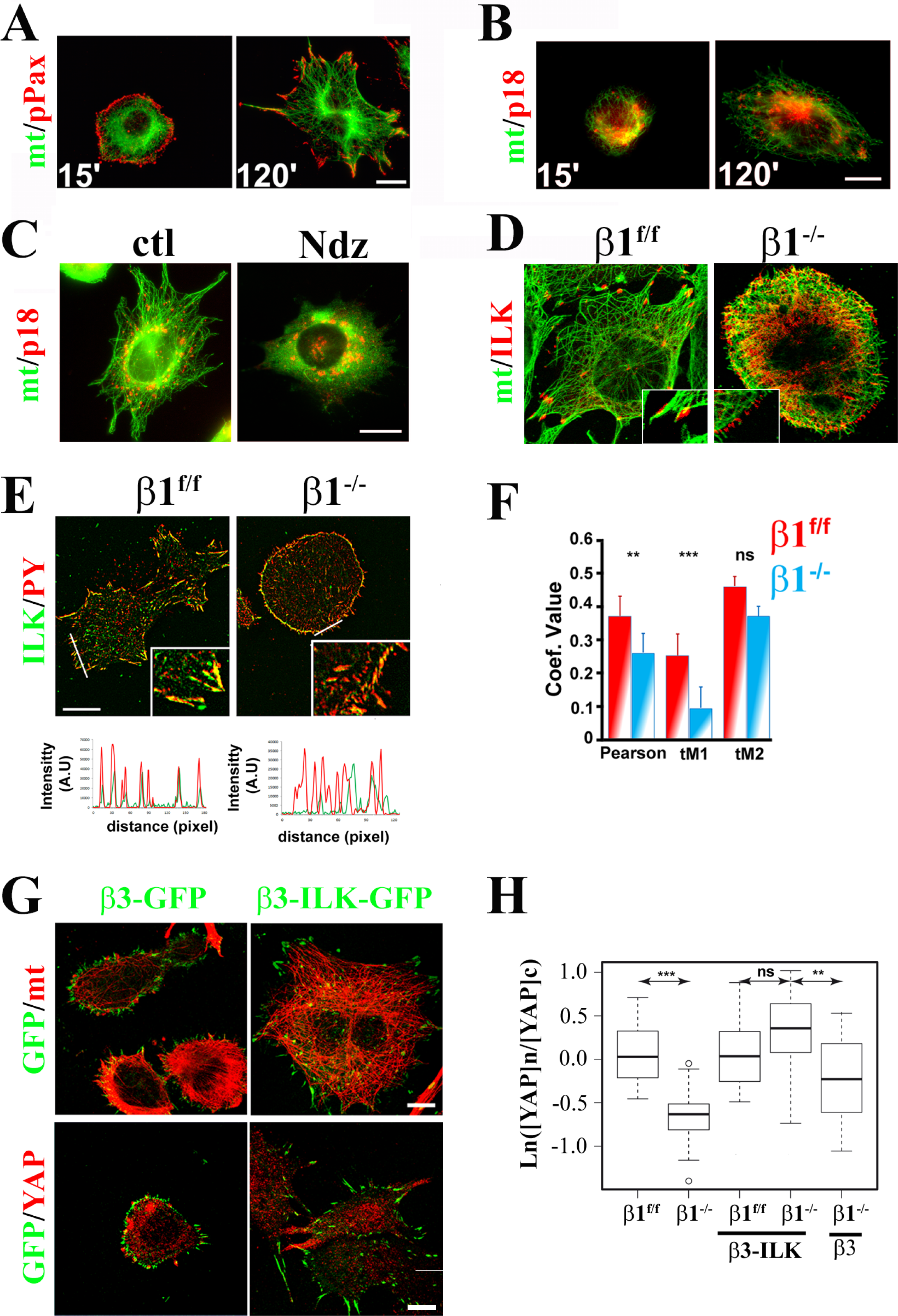
p18/LAMTOR1 distribution and YAP nuclear translocation depend on β1 integrin mediated microtubule anchoring to focal adhesions. **A.** Osteoblast cells were seeded on fibronectin-coated glass coverslips for 15 or 120 min. Microtubule organization was analyzed after tubulin staining and focal adhesion distribution using pPaxillin^Y31^ and anti β-tubulin specific antibodies, respectively. Scale bar represents 10µm. **B.** Osteoblast cells transiently expressing p18-GFP were seeded on fibronectin-coated glass coverslips for 15 or 120 min. p18/LAMTOR1 distribution was imaged (presented in red) together with microtubules (green, stained with β-tubulin). Scale bar represents 10µm. **C.** Osteoblasts expressing p18-GFP were seeded overnight on glass coverslip. p18/LAMTOR1 (red) and microtubules (green) were analyzed in control cells (ctl) and in nocodazole treated cells (Ndz, 10µg/ml, 1h). Scale bar represents 10µm. **D.** β1^f/f^ and β1 deficient osteoblasts stably expressing ILK-GFP (red) were seeded overnight on glass coverslips and stained for microtubules (green). Scale bar represents 10µm. **E.** Control osteoblasts (β1^f/f^) or deficient for β1 integrins stably expressing ILK-GFP were seeded overnight on glass coverslip and focal adhesions stained using an anti phosphotyrosine monoclonal antibody (upper panel). Line intensity profiles were obtained using Icy software plugin. Scale bar represents 10µm. **F.** Histogram representing Pearson’s, thresholded Manders’s coefficients (tM1, tM2) obtained from confocal images of control or β1 integrin deficient osteoblast stably expressing ILK-GFP and stained with anti phosphotyrosine to label focal adhesions. p-value was assessed by a two-tailed unpaired Student’s t-test, with n>20 cells analyzed. **G.** β1 integrin deficient osteoblasts stably expressing either β3-GFP (green, left) or β3-ILK-GFP (green, right) fusion proteins were seeded on glass coverslip and stained for microtubules (red, upper panels) or YAP (red, lower panels). Scale bar represents 10µm. **H.** Statistical analysis of YAP cytoplasmic to nuclear ratio represented in a logarithmic scale. Control cells β1^f/f^ and β1 integrin deficient osteoblast without or stably expressing either β3-GFP or β3-ILK-GFP fusion proteins were seeded overnight on glass coverslips. YAP subcellular localization was then analyzed after indirect immunofluorescence. Intensity values were obtained from confocal images using Fiji software. p-value was assessed by a two-tailed unpaired Student’s t-test, the box plot is representative of 2 independent experiments with n>30 cells analyzed.

It has been reported that focal adhesions, by targeting microtubules in an ILK dependent manner, are required for the polarized delivery of caveolin (a bona fide DRM marker) at the plasma membrane [25]. Indeed, co-staining of the microtubule network with ILK revealed that some microtubules were targeted to focal adhesions supporting these previous data (Fig. 4D, left panel). The loss of β1 integrin expression was characterized by a strong defect in microtubules targeting to focal adhesions (Fig. 4D, right panel). Together, these data nicely supported the idea that β1 integrin dependent establishment of focal adhesions during cell spreading stabilizes microtubules near the plasma membrane in order to allow p18/LAMTOR recycling.

Subsequently, we used β1 deficient cells that display a profound defect in the targeting of microtubules to focal adhesions, to mechanistically unravel the interplay between microtubules, focal adhesions, and YAP signaling. First, ILK recruitment at focal adhesion sites was analyzed in control and β1 deficient cells. Using both transiently and stably expressed ILK-GFP, β1 integrin deficient cells displayed a significant decrease in ILK localization at focal adhesions when compared to control cells (Fig. 4E and Fig. EV4). Indeed, line intensity profile nicely showed a clear localization of ILK at PY positive focal adhesions that was not observed upon β1 integrin removal. This observation was further confirmed by the quantitative analysis of fluorescence colocalization that revealed a significant decrease in both Spearman and thresholded Manders values in β1 deficient cells (Fig. 4F). The decrease of thresholded Manders was significant when the value reported the colocalization extent of ILK (green channel) in focal adhesion (red channel), supporting the reduced recruitment of ILK into focal adhesions. These data suggested a rationale for the need of β1 integrins in targeting of microtubule by controlling ILK recruitment to focal adhesions as previously suggested [30]. To further demonstrate this functional link, a β3-ILK-GFP construct was used to restore ILK localization into focal adhesion in β1^−/−^ cells. As expected, the β3-ILK-GFP integrin subunit was well incorporated into focal adhesions as previously reported [31], and importantly, microtubule targeting to focal adhesions was also restored (Fig. 4G, upper panels). Immunostaining and quantification of cytoplasmic/nucleus YAP ratio revealed a significant increase in YAP nuclear translocation in cells expressing the chimeric β3-ILK-GFP protein compared to β1 deficient cells (Fig. 4G, lower panels and 4H). Collectively these data strongly supported the view that microtubule targeting to focal adhesions (through β1 and ILK) is crucial for YAP nuclear translocation by controlling LAMTOR targeting to focal adhesions.

### p18/LAMTOR1 controls Late Endosomes subcellular distribution

Having demonstrated that the loss of p18/LAMTOR1 negatively impacted on YAP nuclear shuttling, we investigated the molecular basis underlying this process. p18/LAMTOR1 was initially described to be recruited on DRM of late endosomes and to regulate their subcellular distribution [26,32], suggesting that it might be required for proper delivery of signaling proteins associated with late endosomes. While it was demonstrated that the loss of p18/LAMTOR1 or p14/LAMTOR2 affects late endosome distribution, the role of LAMTOR1/2 in controlling late endosome targeting to focal adhesions was not investigated. To this end, GFP-Rab-7 was transfected in sh-scr and sh-p18 cells to follow late endosomes distribution. Indeed, in sharp contrast with control cells that displayed Rab-7 positive vesicles that reached cell periphery, this distribution was seldom observed upon p18/LAMTOR1 silencing (Fig. 5A and 5B). Accordingly, quantification of late endosome targeting to focal adhesions indicated that this process was severely reduced in p18/LAMTOR1 silenced cells when compared to sh-scr control cells (Fig. 5C). This strongly suggested that LAMTOR1 is required for a proper distribution of late-endosomes and in particular for their targeting to focal adhesions. The addition of primaquine, nocodazole, and blebbistatin showed similar results to what was observed for p18/LAMTOR1 distribution, with a reduced density of Rab-7 positive vesicles localizing to the cell periphery (Fig. 5D). Next, we investigated whether Rab-7 distribution also depends on matrix stiffness. Indeed, in line with the p18/LAMTOR1 peripheral distribution that was observed in stiff conditions, Rab-7 distribution was also sensitive to matrix compliance (Fig. 5E). Finally, some Rab-7 vesicles were also targeted to cell edges in a cell adhesion dependent manner, again showing that cell adhesion is a key process in regulating recycling endosomes (Fig. 5F).

**Figure 5.**
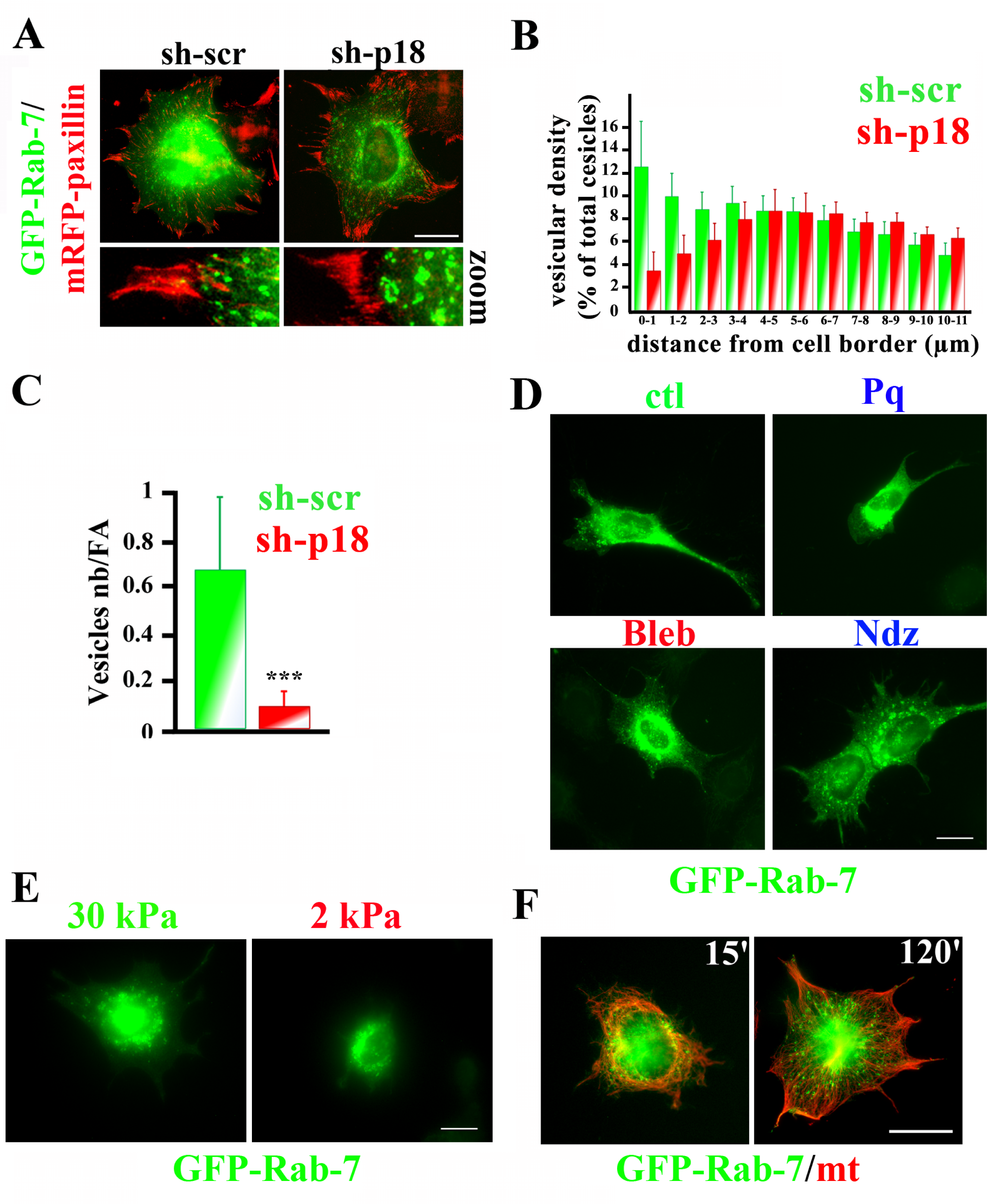
Rab-7 late endosomes subcellular distribution is mechano-dependent and required p18/LAMTOR1. **A.** Sh-scr or sh-18/LAMTOR1 (sh-p18) stably expressing mRFP-Paxillin were transiently transfected with GFP-Rab-7. 24h post transfection cells were seeded on glass coverslip and fixed after overnight. Scale bar represents 10µ **B.** Statistical analysis of Rab7-GFP subcellular distribution in sh-scr or sh-18/LAMTOR1 (sh-p18). GFP fluorescence was imaged and Rab7-GFP distribution was analyzed using Icy software. Histogram represents stepwise particles localization from the cell edges to the cell nucleus displayed as percent of total vesicles. Representative of two independent experiments. p-value was assessed by a two-tailed unpaired Student’s t-test. Differences were not-significant between sh-scr and sh-p18 for 3 to 8; (*) for 8 to 10; (**) for 10 to 12 and (***) for 0 to 3. **C.** Stastistical analysis of GFP-Rab-7 targeting to focal adhesions in control (sh-scr, green) and sh-p18/LAMTOR1 (sh-p18, red). p-value was assessed by a two-tailed unpaired Student’s t-test, with n = 20 cells analyzed. **D.** Subcellular localization of GFP-Rab-7. Control osteoblasts expressing GFP-Rab-7 were treated or not with blebbistatin (Bleb, 1h, 10µg/ml), primaquine (Pq, 1h, 50µg/ml) or nocodazole (Ndz, 10µg/ml, 1h). **E.** Subcellular localization of GFP-Rab-7 in osteoblast cells spread for 2h on fibronectin-coated PDMS hydrogel of different stiffness (Young moduli are 30 kPa, 2kPa). Scale bar represents 10µm. **F.** Osteoblast transiently expessing GFP-Rab-7 were seeded on fibronectin coated glass coverslip for 15 or 120 minutes. Cells were fixed and microtubules network visualized after staining β-tubulin. **G.** **H.** Histogram representing Pearson’s coefficient, thresholded Manders’s green in red (tM1) and thresholded Manders’s red in green (tM1) obtained from confocal images of control osteoblast cells or β1 integrin deficient cells stably expressing ILK-GFP and stained with phosphotyrosine specific antibodies to label focal adhesions. Images were analyzed using the Fiji Jacop plugin. p-value was assessed by a two-tailed unpaired Student’s t-test, with n>20 cells analyzed. **I.** Statistical analysis of Rab7-GFP subcellular distribution. Osteoblasts expressing either a sh-scr (red), sh-p18/LAMTOR1 (green) were transfected with GFP-Rab7 and imaged for GFP fluorescence. Vesicular distribution was analyzed using Icy software. Histogram represents stepwise particles localization from the cell edges to the cell nucleus displayed as percent of total vesicles. p-value was assessed by a two-tailed unpaired Student’s t-test, with n>20 cells analyzed. **J.** Statistical analysis of GFP-Rab-7 targeting to focal adhesions in control (sh-scr, red), and sh-Rab-11 expressing cells (green). p-value was assessed by a two-tailed unpaired Student’s t-test, with n>20 cells analyzed. **K.** β1^f/f^ osteoblast were transfected with GFP-Rab7 and 24h post-transfection, the cells were seeded on glass coverslips. After overnight incubation the cells were treated for 1h with blebbistatin (Bleb, 10µg/ml), primaquine (Pq, 50µg/ml), nocodazole (Ndz, 10 µg/ml) or seeded on fibronectin coated PDMS hydrogel for 2 h. (2kPa and 30 kPa). Scale bar represents 10µm. **L.** β1^f/f^ osteoblasts were transfected with GFP-Rab7 and 48h post-transfection the cells were seeded on fibronectin coated glass coverslips for 15 min and 120 min and microtubules stained using anti tubulin antibodies. Scale bar represents 10µm.

### p18/LAMTOR1 regulates Src signaling

Altogether, the above presented data suggested that the recycling of late endosomes is required for YAP nuclear translocation, likely by regulating the local delivery of signaling proteins. Among the strongest candidate, the non-receptor Src tyrosine kinase was identified to be an important regulator of YAP nuclear translocation and Src is also found to traffic back to the plasma membrane [14,33,34]. Therefore, we hypothesized that Src could recycle back to the plasma membrane through p18/LAMTOR1 positive vesicles. To verify this assumption, we first asked whether Src could be colocalized with p18/LAMTOR1 positive vesicles. Since, the use of antibodies for detecting endogenous Src was giving inconsistent results likely due to the relatively low level of Src expression, we transiently co-expressed fluorescently tagged Src (mCherry-Src) and p18/LAMTOR1 (p18-GFP) in control cells. Confocal imaging revealed that mCherry-Src was not only localized diffusely within the cytoplasm but also appeared as punctuate staining in good agreement with previously report [35]. These vesicular-like structures were also positive for p18/LAMTOR1 (Fig. 6A). These observations were further supported by colocalization analysis that revealed a significant Spearman coefficient above 0.5 and a thresholded Manders (tM1) values up to 0.7 when the green channel (p18/LAMTOR) overlapping with the red channel (mCherry-Src) was analyzed. The smaller tM2 value (red overlapping with the green) likely reflects the large diffusible fraction of Src (Fig. 6B). Finally, we asked whether Src and p18 were dynamically coupled. To address this question, mCherry-Src and p18-GFP were transiently transfected in SYF cells (Src, Yes and Fyn triple knockout) to avoid any contribution of other major Src family members. Indeed, time lapse videomicroscopy further supported previous data showing a tight connection between Src and p18/LAMTOR1 (Movie EV5). All together these data strongly suggested that a pool of Src colocalizes with p18/LAMTOR1. Since we previously reported that p18/LAMTOR1 controls late endosomes distribution we wondered whether Src distribution was also p18/LAMTOR1 dependent. Src distribution was then analyzed in both p18/LAMTOR1 knocked-down cells (sh-p18) and compared to control cells (sh-scramble) and late endosomes visualized using Rab-7-GFP. In line with the putative localization of Src on late endosomes, we observed that Src and Rab-7 were colocalized in control cells as visualized by an extensive yellow signal in the merge image (Fig. 6C, left panels and blue insert). However, in sh-p18 cells and when peripheral vesicles were specifically analyzed we noticed a significant reduction in the colocalization between both proteins (Fig 6C, right panels and blue insert). Interestingly, this defect was not less obvious on vesicles distant from the periphery suggesting that p18/LAMTOR1 may not be required for Src recruitment on late endosomes but rather for its recycling to the plasma membrane (Fig. 6C, red inserts). This colocalization between Src and Rab-7 was further confirmed by digital quantification of images and with the resulting decrease of both the Spearman and Manders coefficient (Fig. 6D).

**Figure 6.**
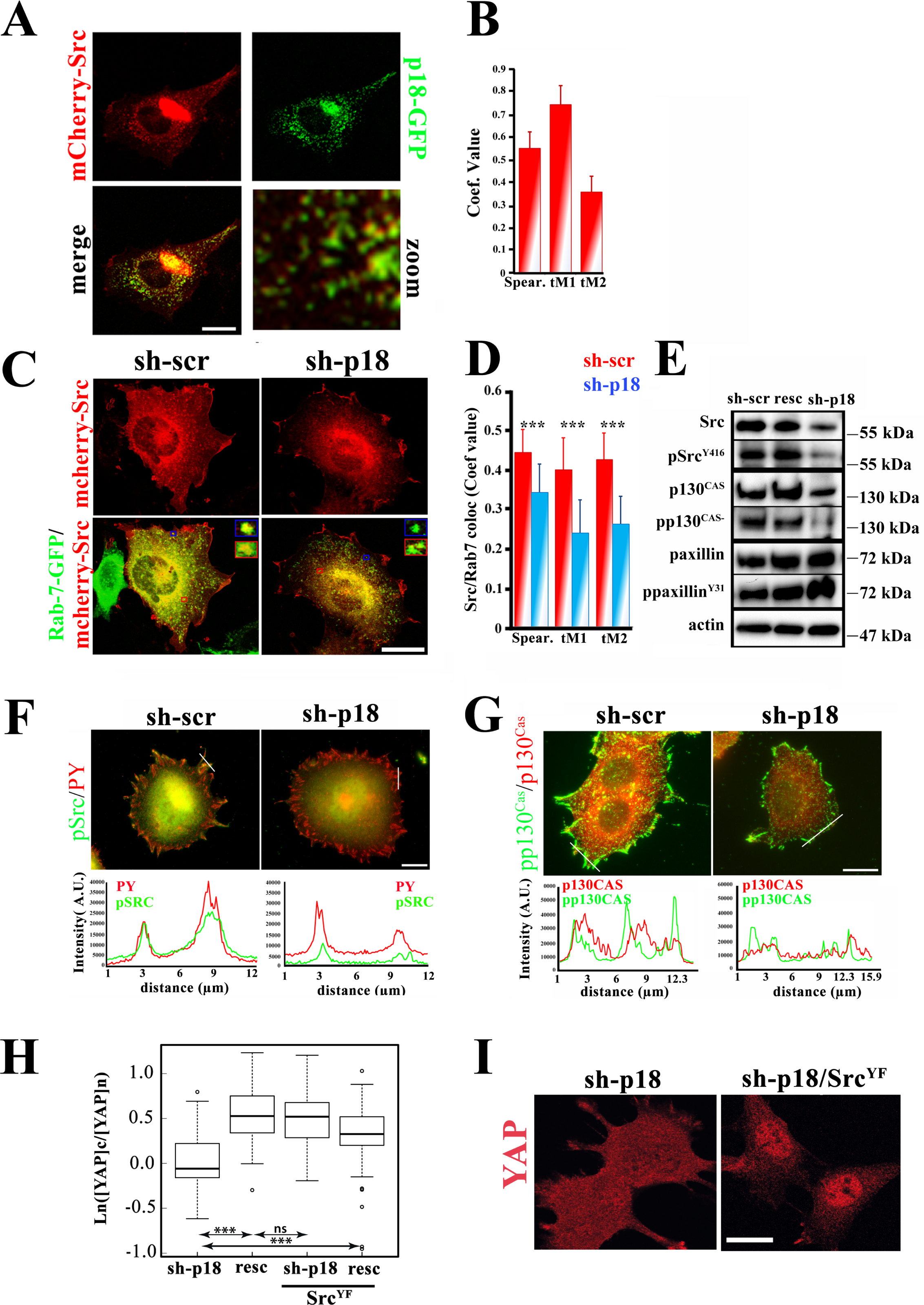
p18/LAMTOR dependent Src delivery to the plasma membrane controls YAP nuclear shuttling. **A.** Control osteoblast cells were transiently transfected with p18-GFP and mCherry-Src. 24h post-transfection the cells were seeded on glass coverslip and GFP and mCherry channels acquired using confocal microscope. Scale bar represents 10µm. **B.** Histogram representing Pearson’s coefficient, thresholded Manders’s (tM1, tM2) obtained from confocal images of p18-GFP and mCherry-Src cotransfected osteoblast cells. Images were analyzed using the Fiji Jacop plugin. p-value was assessed by a two-tailed unpaired Student’s t-test, with n>35 cells analyzed. **C.** Control osteoblast cells (sh-scr) or sh-p18/LAMTOR1 (sh-p18) were transiently transfected with GFP-Rab-7 and mCherry-Src. 24h post-transfection the cells were seeded on glass coverslip and GFP and mCherry channels acquired using confocal microscope. Scale bar represents 10µm. **D.** Histogram representing Pearson’s coefficient, thresholded Manders’s (tM1, tM2) obtained from confocal images of GFP-Rab-7 and mCherry-Src cotransfected in control (red, sh-scr) or sh-p18/LAMTOR1 (blue, sh-p18) osteoblast cells. Images were analyzed using the Fiji Jacop plugin. p-value was assessed by a two-tailed unpaired Student’s t-test, with n>35 cells analyzed. **E.** Western blot analysis of Src, p Src^Y146^, p130^CAS^, p-p130^CAS^, paxillin, paxillin^Y31^ form lysate of control cells (sh-scr), p18/LAMTOR1 silenced cells with (resc) or without (sh-p18) p18/LAMTOR1-GFP expression. Actin was used as internal loading control. Representative of 3 independent experiments. **F.** Control cells (sh-scr) and p18/LAMTOR1 silenced cells (sh-p18) were seeded overnight on glass coverslips and stained with pSrc^Y416^ to detect the active form of Src and phospho tyrosines to label focal adhesions. Line intensity profiles were obtained using Icy software plugin. Scale bar represents 10µm **G.** Control cells (sh-scr) and p18/LAMTOR1 silenced cells (sh-p18) were seeded overnight on glass coverslips and stained with p-p130^CAS^ and p-130^CAS^. Line intensity profiles were obtained using Icy software plugin. Scale bar represents 10µm **H.** Statistical analysis of YAP cytoplasmic to nuclear ratio represented in a logarithmic scale. Sh-p18/LAMTOR1 cells expressing the active form of Src (Src^YF^) or not (sh-p18) and rescued cells cells expressing the active form of Src (Resc-Src^YF^) or not (Resc) were seeded overnight on glass coverslips. YAP subcellular localization was then analyzed after indirect immunofluorescence. Intensity value was obtained from confocal images using Fiji software. p-value was assessed by a two-tailed unpaired Student’s t-test, the box plot is representative of 2 independent experiments with n>30 cells analyzed. **I.** Subcellular localization of YAP in osteoblast cells expressing stable sh-p18/LAMTOR1 with (sh-p18 Src^YF^) or without (sh-p18) the active form of Src^YF^. Scale bar represents 10µm.

Next, we investigated whether p18/LAMTOR1 could control Src activity and the phosphorylation of well-known downstream targets such as p130^CAS^ (Fig. 6E) or paxillin (Fig EV5). Src activity, as reported by Western-blotting of the phosphorylated form of Src (Y416), revealed a strong diminution of activated Src in p18 silenced cells compared to control and rescued cells. It is also noteworthy that the total amount of Src was severely reduced in sh-p18 cells likely explaining the reduced level of activated Src (Fig. 6E). When the subcellular distribution of phosphorylated Src was investigated we observed an almost lack of pSrc that normally localized at cell edges (Fig. 6F). Accordingly, with this reduced level of activated Src in p18 silenced cells, the well-recognized Src substrate p130^CAS^ followed a parallel decrease in phosphorylation (Fig 6E). While the phosphorylation of p130^CAS^ was mainly observed at focal adhesion sites in control cells, at these locations this phosphorylation was significantly reduced upon silencing of p18/LAMTOR1 (Fig. 6G). However, when paxillin phosphorylation was investigated, silencing of p18/LAMTOR resulted in no significant difference compared to controls (scr and rescued; Fig. 6E). At the subcellular level, paxillin was extensively phosphorylated in both sh-p18 cells and control cells (Fig. EV5).

Together these data pinpointed an important role for LAMTOR in regulating Src signaling at the cell periphery by controlling late endosomes distribution. Finally, to directly link the defect in Src activity to YAP nuclear translocation, sh-p18 cells were transduced with the activated form of Src (Src^Y527F^) and YAP subcellular localization analyzed and quantified (Fig. 6H and 6I). The expression of activated Src led to the relocalization of YAP into the nucleus showing that forcing Src activity bypassed the requirement of vesicular transport. Altogether, these results propose that p18/LAMTOR1 expressed on recycling endosomes is required for Src trafficking back to the plasma membrane in order to drive YAP nuclear translocation.

## Discussion

Here we present evidence for a role of vesicular recycling in the regulation of YAP nuclear shuttling. Our work adds to several reports proposing vesicular trafficking as an integrating process regulating important cell signaling pathways and thereby cell behavior such as proliferation, spreading or migration [6,36,37]. Indeed, we uncovered that the nuclear translocation of YAP, a well-known co-transcription factor involved in cell proliferation, required an active vesicular trafficking back to the plasma membrane. Mechanistically, β1 integrin dependent cell adhesion, by allowing anchoring (through ILK) of microtubules to focal adhesion sites, permits the recycling of p18/LAMTOR1 positive vesicles for local delivery of Src to the plasma membrane - a necessary mechanism for YAP nuclear translocation.

It is well known that anchorage-dependent cell growth requires a firm cell adhesion to the extracellular matrix, a process in which YAP signaling is likely playing a significant role [1,24,28]. Our work extends these data, and proposes that the recycling of late endosomes to the plasma membrane is equally required for the targeting of the non-receptor tyrosine kinase Src to the plasma membrane to promote YAP nuclear translocation. These findings support the emerging role of Src in regulating YAP nuclear translocation downstream of cell adhesion [13,14,34,38]. The exact role of Src in this regulation is still under debate and different alternative pathways have been proposed. On the one hand, Src was shown to directly phosphorylate YAP or its upstream kinase LATS1/2 and thereby promotes YAP nuclear translocation [39,40]. One the other hand, Src is a well-known regulator of Rac-1 and this later protein was described being involved in YAP nuclear translocation as well [14,41]. Several lines of evidences are supportive of this later view. Firstly, v-Src induced cell transformation relies on the PAK1/Rac-1 binding protein β-Pix/Cool1 that was also shown to induce Rac-1 recruitment at membrane ruffles [42–44]. Secondly, β-Pix/Cool1 was identified in a high throughput screening as a regulator of YAP nuclear translocation [46]. Supporting this hypothesis, lowering Src activity either upon p18/LAMTOR1 silencing or pharmacological inhibition led to a reduced localization of Rac1 at protrusive borders (Fig. EV6). It is noteworthy that the phosphorylation of merlin by the Rac-1 effector PAK-1 was reported to inhibit its interaction with both LATS and YAP, thereby promoting YAP dephosphorylation and nuclear translocation [14]. Therefore, this pathway appears to be an alternative to a direct role of Src on YAP and LATS. It will be important in the future to investigate the relative contribution of those specific pathways in the control of YAP activity.

YAP nuclear translocation is actively mediated by inputs coming from the extracellular matrix such as stiffness and chemical composition. As expected, integrins, as the main receptors that integrate those extracellular cues such as contractility into cellular signaling, appear to play a critical role in this process. Our data support this view and are consistent with previous reports showing that both β1 integrins and ILK are key players regulating YAP signaling [23,46]. Despite the fact that the involvement of ILK in the control of YAP activity has been demonstrated, the molecular mechanism involved remains puzzling. Indeed, from the above-mentioned work it was proposed that ILK regulates YAP nuclear translocation by phosphorylating and inhibiting the merlin phosphatase MYPT-PP1. However, the kinase activity of ILK is highly disputed by several groups and scaffolding function for ILK has been alternatively proposed [48]. Although we cannot exclude the involvement of the ILK kinase activity in YAP nuclear translocation, our data would favor its scaffolding function by supporting microtubule anchoring to focal adhesions as previously reported [4,47]. Whatever the exact mechanism is, it appears that β1 integrins, by regulating the recruitment of focal adhesion proteins such as ILK, has a major role in integrating extracellular cues to YAP nuclear shuttling.

This relationship between integrins, YAP signaling, and the extracellular environment provides a rational framework for the progression of solid tumors. YAP signaling is well-known to be regulated by cell contractility and stiffness, however, whether external cues affect vesicular trafficking has much less attracted attention. In the same line, caveolin was recently reported to be involved in YAP response to extracellular stiffness [45]. Interestingly, caveolin is involved in DRM trafficking and was proposed to promote Rac-1 withdrawal from the plasma membrane upon cell detachment or on low matrix stiffness [7,24,48]. Hence, vesicular trafficking either via controlling Rac-1 internalization or its accumulation via the recycling of late endosomes appears to be a potent way to control YAP nuclear shuttling and, thereby, anchorage-dependent growth.

Our data also add to the growing evidence identifying late endosomes as an important cell signaling compartment [26,49]. Their signaling function appears to be tightly correlated to their dynamics and more specifically to their capacity to recycle back to the plasma membrane near or at focal adhesion sites [50,51]. Isolated from late endosomal DRMs, the LAMTOR complex is involved in the regulation of their dynamics and signaling capabilities [26,32]. In good agreement with these reports, we observed a critical role for p18/LAMTOR1 in the targeting of late endosomes to focal adhesion sites. We further showed that Src is traveling via this late endosomal compartment. Interestingly, Src lipid modification confers its specific subcellular dynamic over that of other family members such as Fyn and Yes [35]. While it was known that Src recycles back to the plasma membrane, the nature of the recycling vesicles has been disputed [52,53]. Here we showed that late endosome recycling requires an active vesicular trafficking (Rab-11 dependent and primaquine sensitive) and a functional microtubule network anchored to focal adhesions. The function of those late endosome at focal adhesions is still poorly understood. It was previously reported that the docking of late endosomes to focal adhesion regulates their turn-over [29]. We uncovered another function for those vesicles in the regulation of YAP nuclear translocation. Moreover, considering the role of Src in focal adhesion turnover [54,55], our data also opened up the possibility that the reduced focal adhesion dynamics reported in LAMTOR2 deficient cells might also results from a lack of Src activity near focal adhesions. It is noteworthy that LAMTOR, through p14, interacts with IQGAP-1, a direct partner of both Rac-1 and the exocyst complex [29,56,57]. Definitely more investigation will be required in the future to fully address this connection between focal adhesions, late endosomes and the exocytosis that appear to be involved the mechano-dependent integration of extracellular inputs.

## ACKNOWLEDGMENTS

We would like to thank Drs Manié, Zerial, Roche, Van Obberghen, Okada, Gertler, Fässler, McCaffrey, Hiraishi and Gauthier-Rouviere for sharing their tools and reagents. Ms Shalini Chandrashekhar for reading and editing the manuscript. Mr Mazzega for its technical assistance with cell imaging (Cellular imaging was performed at the Optical Microscopy -Cell Imaging (Microcell) core facility of the Institute of Advanced Biosciences). This work was supported by a grant from the SFCE (CRAUFESD16), the Finovi foundation and ANR (15-CE14-0010-03).

## AUTHOR CONTRIBUTIONS

Conceptualization: DB; Methodology: MRB, MB, MP and DB; Investigation: MRB, MB, TZ, GC, DB; Writing: MRB, MB, BWH, PR, DL and DB; Supervision: DB and MRB.

## Materials and Methods

### Cell lines

β1^f/f^ and β1^−/−^ cell line generation and characterization was presented previously [14]. From these original cell lines, sh-p18 (Santa Cruz Biotechnology sc-36146-v), sh-Rab11a (addgene # 26710, Dr K. Mostov) and sh-srcamble (addgene #17920, Dr S. Stewart) were generated by trandsuction of lentivirus particles. Cells were maintened on puromycin selection. All other cell lines were generated upon retrovirus transduction and transgene expression was verified by Western blotting and/or immunostaining.

### Antibodies and expression vectors

Anti-YAP^S127^, anti-Rab-11, anti-Src, anti-Src^Y416^ anti-p130^CASY410^, was from Cell Signaling (Ozyme, St Quentin en Yvelines France). Anti-YAP, anti-β-tubulin (clone 2.1) was from Santa Cruz (Heidelberg, Germany). Mouse β1 integrin (MB1.2), mouse/human β1 integrin (9EG7) and anti-Rac1, p130^CAS^ were from BD Biosciences (Le Pont de Claix, France). Anti actin, was from Sigma Aldrich (L’Isle d’Abeau France). Anti-LAMP1, anti-p-PAK was from abcam. Anti-paxillinY31 was from Invitrogen. Anti-Paxillin was from Millipore. The anti-phosphotyrosine monoclonal antibody 4G10 used as hybridoma supernatant was produced in our laboratory. Rabbit polyclonal antibodies against p18/LAMTOR1 were a generous gift from Dr. S. Manié (CRCL, Lyon, France).

The human β1-expressing construct was based on the pCL-MFG retroviral vector as described previously [58]. pCL-MFGβ3-GFP-ILK and pCL-MFG-β3-GFP and pCL-MFG-hILK-EGFP were a gift from Drs E. Van Obberghen and R. Fässler (iBV, Nice, France; MPI, Martinsried, Germany, respectively). pBabe-Neo-Src^YF^ was from Pr. M. Humphries (Manchester University, UK). pBABE-puro-FlagYAP2 was from Dr. M. Sudol (Addgene #27472). Flag tagged YAP2^5SA^ was from Dr. K.L. Guan (Addgene #27371). mCherryYAP constructs were engineered from these initial plasmids and cloned into pCL-MFG retroviral vectors. The pEGFP-Rac1^G12V^ plasmid was a gift from Dr C. Gauthier-Rouvière (CRBM, Montpellier, France). The insert GFP-Rac1^G12V^ was subcloned into the retroviral vector pBaba-puro. Wild type and dominant negative mutants of Rab proteins were obtained from Dr M. McCaffrey. Human wild-type (WT) paxillin cDNAs were subcloned from pBABE vectors generously provided by Dr M. Hiraishi (Osaka Bioscience Institute, Osaka, Japan), mRFP-paxillin vector was engineered from this initial vector. pEGFP-N1-p18 was an original gift from Dr Masado Okada (Osaka University, Japan) and was subcloned into pCLM-FG retroviral vectors. The pmCherry-N1-Src was from Dr S. Roche (CRBM, Montpellier, France). Dynasore, Blebbistatin, Nocodazole and Primaquine were from Sigma Aldrich.

### Transfections and Infections

HEK GP 293 cells (Clontech) were transfected with plasmid DNA using TurboFect Transfection reagent (ThermoFisher scientific, Courtabeuf, France) according to manufacturer’s instructions. Osteoblast retroviral infections were performed as previously described [59].

### Quantification of Microscopy

Cells grown on glass coverslips were fixed with 4% paraformaldehyde and 5% sucrose in PBS for 10 minutes at room temperature (RT), then permeabilized in 0.1% Triton X-100 in PBS, for 5 minutes. Coverslips were washed twice with PBS, blocked in 1% BSA in PBS and when needed incubated for 1 hour at RT with primary antibodies. Cells were rinsed in PBS and secondary antibodies were added for 1 hour at RT. Coverslips were permanently mounted in Mowiol from Calbiochem (VWR International, Strasbourg, France) containing 4’6-diamidino-2-phenylindole (DAPI). Fixed cells were examined using a confocal laser-scanning microscope (LSM 510, Carl Zeiss, Jena, Germany), equipped with a Plan Apochromat 63X oil-immersion objective, N.A.1.4. The pinhole was adjusted to one Airy unit.

### Quantification of YAP nuclear localization

Cells were immuno-stained with an anti YAP and immuno-microscopy was carried out with a confocal laser scanning microscope (LSM510, Carl Zeiss, Jena, Germany) equipped with a 63X plan-Apochromat oil immersion objective (N.A. 1.4) and a pinhole set to one Airy unit. On each cell image a ROI was defined positioned either within the nucleus, or in the cytoplasmic area next to the nucleus envelope. Since the thickness of the two ROI positions were likely identical, the average fluorescence intensity is likely proportional to YAP concentration and was estimated using Image J (NIH Image Software). Within the same cell, the ratio of both fluorescence intensities reflects YAP concentration ratio in both compartments. This ratio was represented under a logarithmic scale in order to have an identical range for positive and negative ratios. Measurements were performed with n ≥ 50 (unless indicated otherwise) and statistical significance was estimated with Student’s test. Boxplots were performed using R public software.

### Videomicroscopy and TIRF

For live imaging, cells were seeded at subconfluent densities in labtek chambers and allowed to grow overnight prior to imaging in DMEM supplemented with 10% FCS, placed on a 37°C heated stage in 5% CO_2_ atmosphere, and imaged with the same Zeiss Axiovert 200M microscope (Carl Zeiss Microscopy, Jena, Germany) equipped with CoolSNAP HQ2 camera (Photometrics, Tucson, USA), 100X (NA 1.46) Plan-Apochromat objective and filters sets to specifically detect Alexa488/GFP or Alexa546/pTRFP. Time lapse was 5 sec interval. Total internal reflection fluorescence (TIRF) microscopy was carried out with the same set-up equipped with the TIRF 2 slider (Carl Zeiss Microscopy, Jena, Germany).

### Vesicular distribution

Cells were imaged using an Axioimager Z.1 microscope equipped with a 63x/1.4 Plan-Apochromat oil objective and an Axiocam Mrm CCD camera controled by Axiovision software (Carl Zeiss Microscopy, Jena, Germany). Images were then analyzed using the Icy software (http://icy.bioimageanalysis.org) with graphical programming tools for protocol editing. First, channels were separated and processed to automatically detect cells borders with the best threshold, HK-means and active contours plugins. In parallel, vesicles were detected using the wavelet spot detector block and the distance between vesicles cell border quantified using the ROI inclusion analysis plugin. Detailed protocol is provided upon request.

### PDMS hydrogels

PDMS hydrogel of different stifness (2, 10 and 30 kPa) were obtained from Excellness Biotech SA (Lausanne, Swizterland). Hydrogels were coated with bovine plasma fibroectin (2.5 µg/ml) according to the company protocol. Cells were seeded for 2-3 hours at 37°C in CO2 and humidified incubator. After washing cells were PFA (3% with 4% sucrose) fixed for 15 minutes and then imaged as previously described.

### Colocalization analysis

Images were obtained with a confocal laser scanning microscope (LSM510-META, Carl Zeiss, Jena, Germany) equipped with a Plan-Apochromat 63x/1,4 oil objective. Optical sectioning was set to 0.8µm. LSM images were then analyzed using Fiji software (NIH software) to run the JACOP plugin. Except when indicated the whole cells was considered as ROI for analysis.

### Quantification of vesicles targeting

Cells were imaged using an Axioimager Z.1 (Carl Zeiss Microimaging) microscope equipped with a 63x/1.4 plan-Apochromat oil objective and an Axiocam Mrm CCD camera controled by Axiovision software (Carl Zeiss Microscopy, Jena, Germany). Images were then processed using Fiji software, first a substract background with a rolling ball of 50 was applied on p18-GFP channel and then mRFP-VASP channel was merged and vesicles in direct contact with focal adhesion (FA) were manually counted using the counter plugin. Then the mean value of attached vesicles was divided by the number of FA within the cell to obtain the vesicle/FA ratio. 20 cells were analyzed for each condition.

### Statistics

Cell analyses were carried out with a minimal cell number of 20 per experimental condition (for most of the presented experiments n was ranging between 50 and 100). Statistical relevance was estimated with Student’s test with bilateral distribution and unequal variance. A p value below 0.01 was considered as significant.

---

**Fig EV1.**
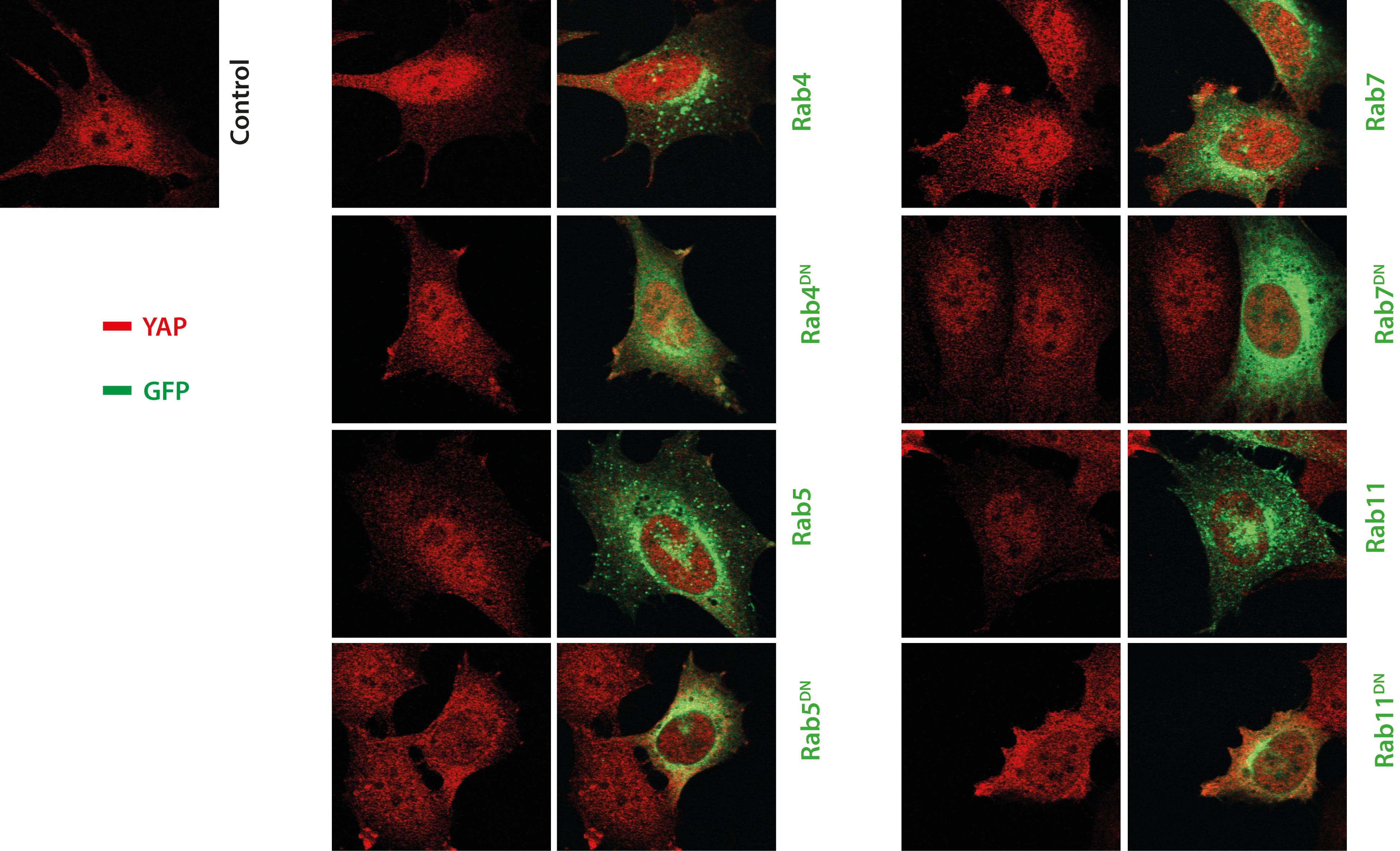

**Fig EV2.**
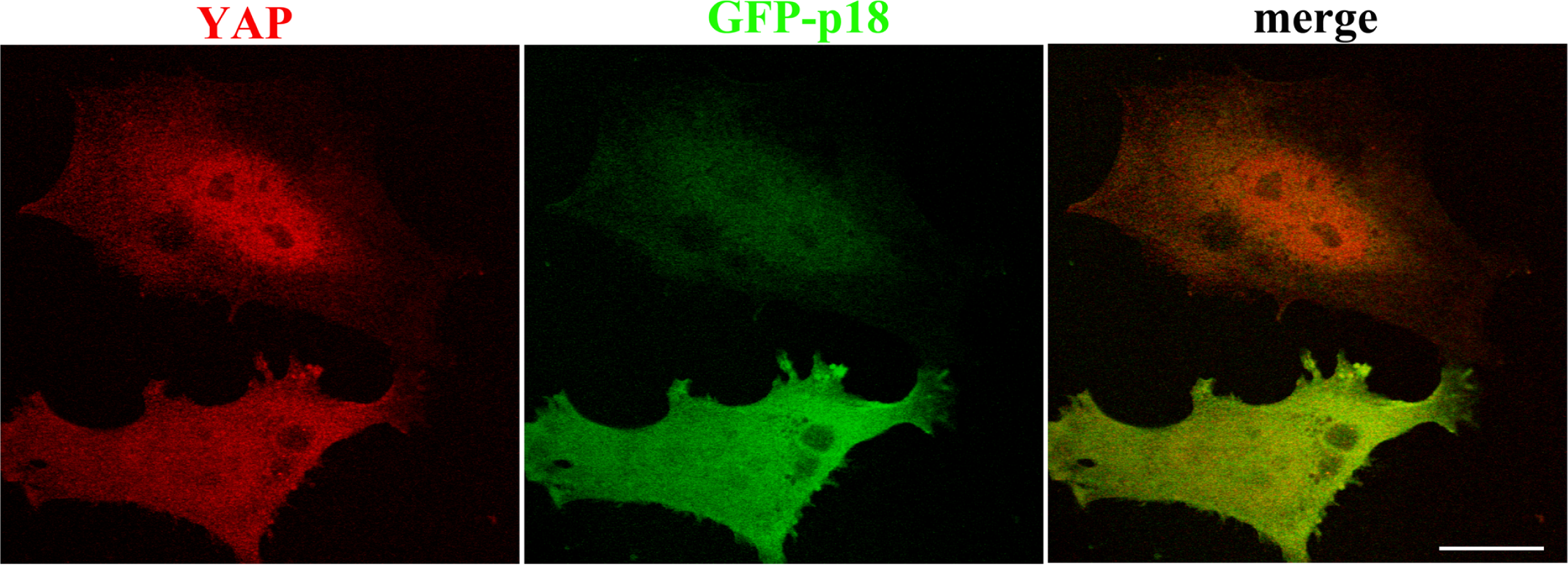

**Fig EV3.**
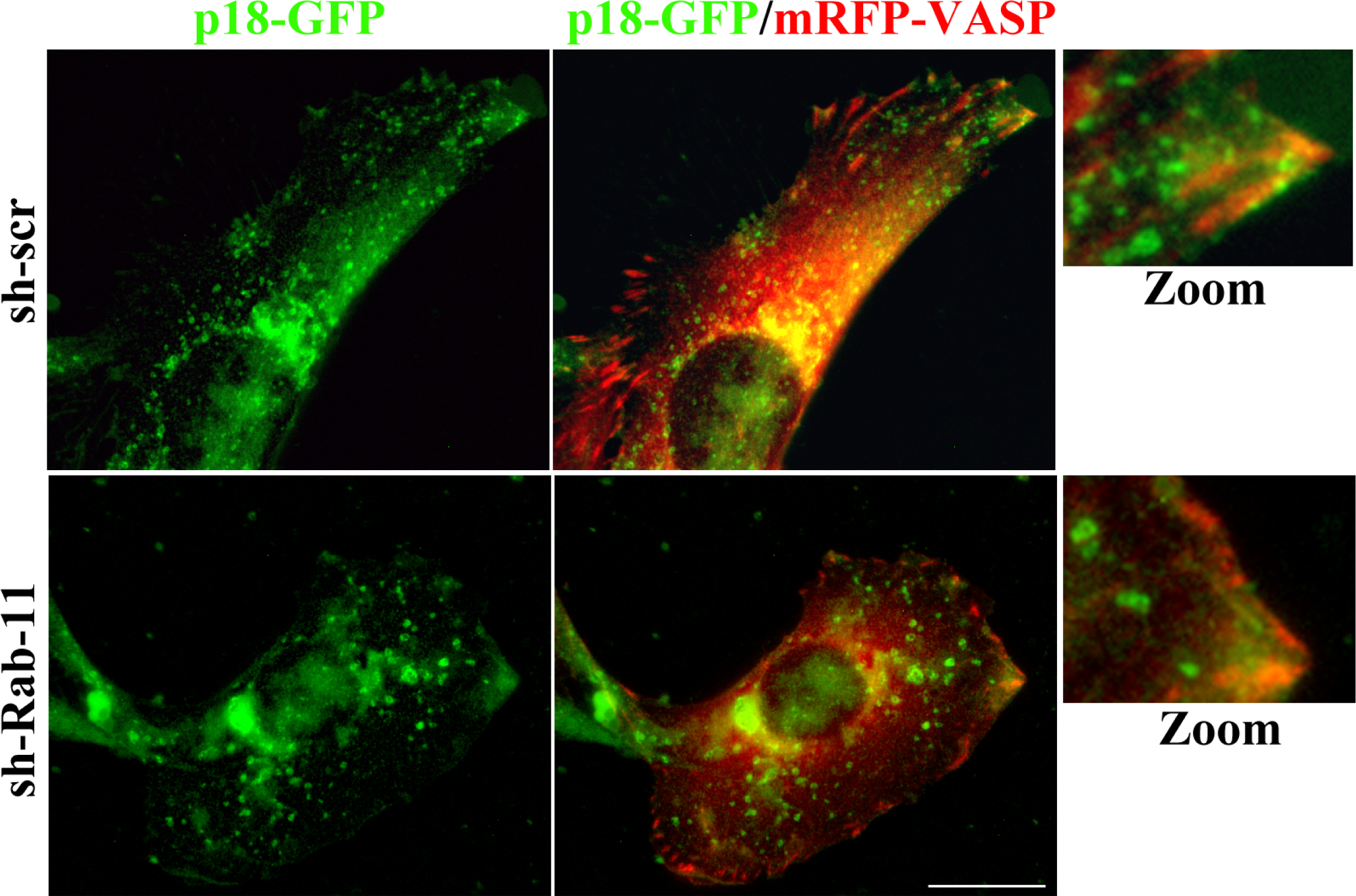

**Fig EV4.**
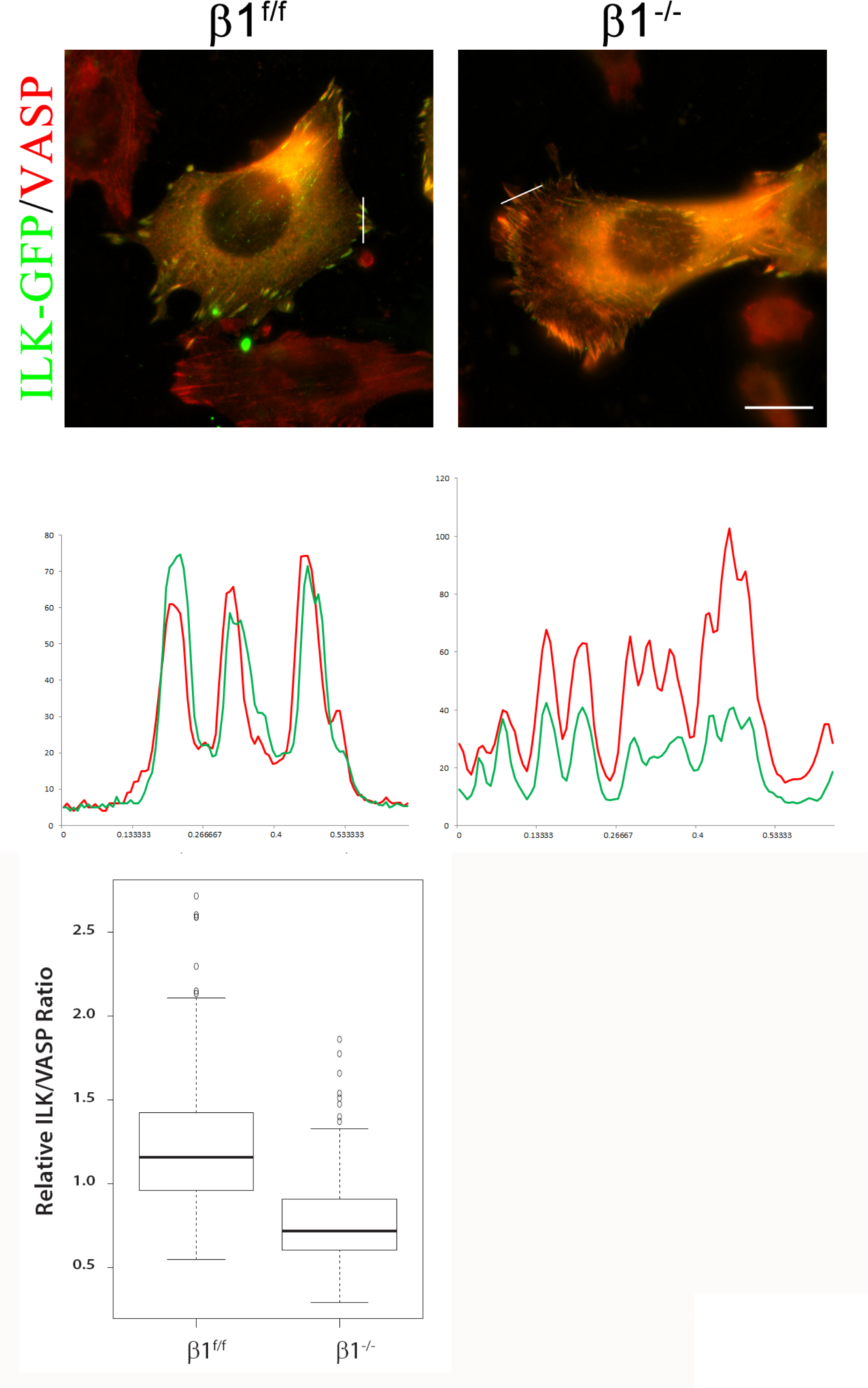

**Fig EV5.**
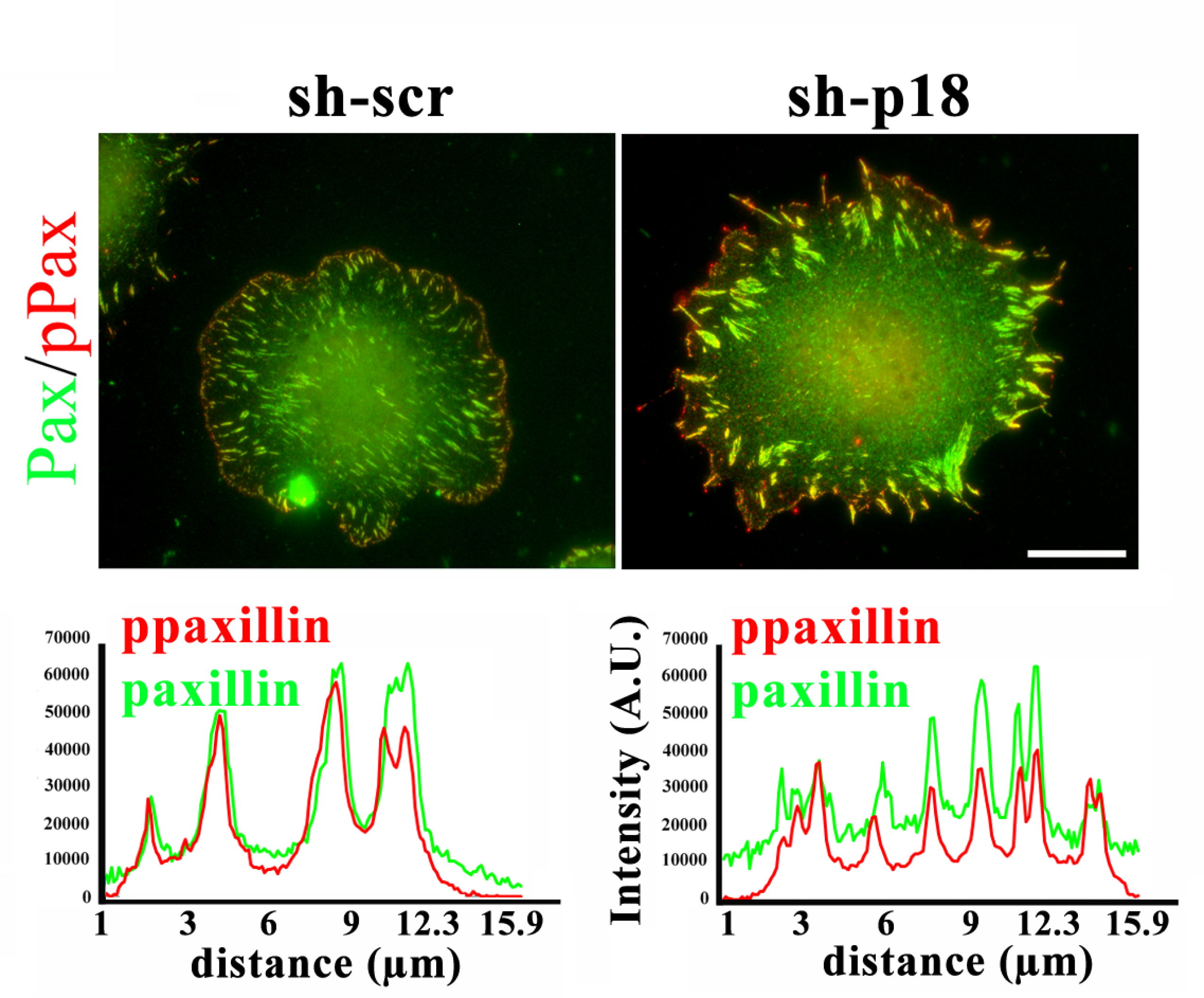

**Fig EV6.**
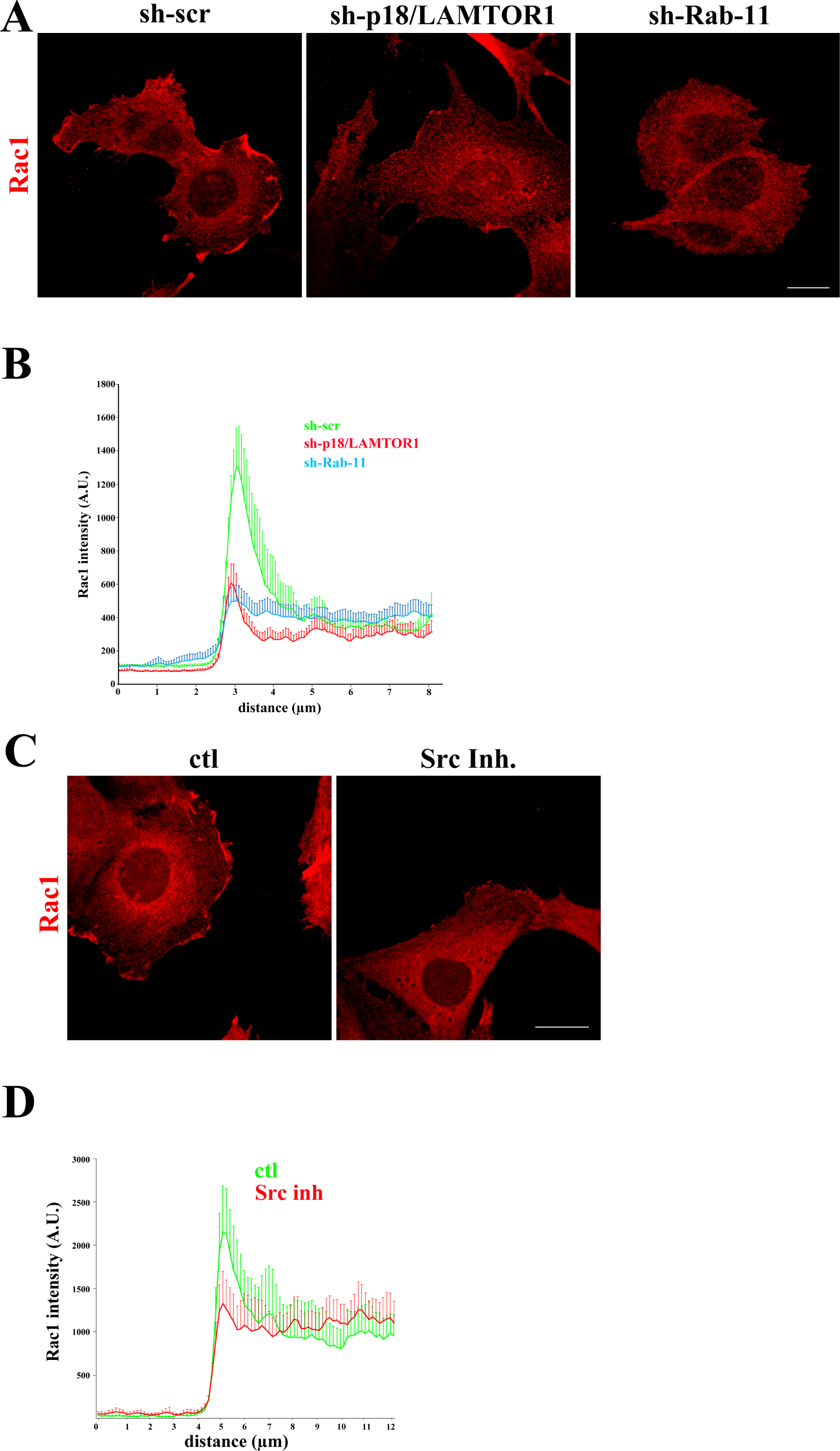

